# Active non-redundancy and viral orchestration sustain diel microbial successions in the coastal ocean

**DOI:** 10.64898/2026.06.22.733315

**Authors:** Olivier Pereira, Elizabeth A. Ottesen, Amber N. DePoy, Pierre Ramond, Songze Chen, Fanru Ji, Shengwei Hou, Nianzhi Jiao, Chuanlun Zhang

## Abstract

Ecosystem resilience in dynamic coastal oceans is conventionally ascribed to functional redundancy, where taxonomically distinct microbes buffer environmental fluctuations through interchangeable metabolic roles. In this study, we reveal a deterministic succession architecture sustained by active non redundancy and temporal metabolic coupling, uncovered via autonomous drone array metatranscriptomic sampling in Daya Bay, China. By resolving the transcriptional landscape into six chronometrically phased modules rather than simple time points, we demonstrate a near total renewal of the active gene pool, with greater than 90% of transcribed clusters showing phase exclusivity within approximatively a two-hour window. This radical functional reshaping exposes a tight, scale dependent taxonomic functional coupling where community composition is a strong, linear predictor of metabolic output, peaking at the genus level (*p* < 0.0001). Functional continuity is maintained not by redundant generalists, but by a precisely sequential relay of specialists, each optimized for transient microniches with minimal overlap between successive phases. Deep learning structural adjudication of the *psbA* gene pool further reveals viral orchestration of this temporal coupling, where incoming cyanophages introduce structurally distinct protein variants that sustain photosynthetic electron flow under peak irradiance. These findings redefine coastal microbiome stability as a fine-tuned rapid succession of temporal specialists rather than a redundant backdrop. We propose a fundamental revision of marine ecosystem models, shifting from passive buffering frameworks to deterministic, clock driven architectures, which may prove to be critical for forecasting microbiome responses to accelerating short term climate variability.

## Introduction

Marine microbial communities represent the dominant fraction of oceanic biomass and act as the principal engines of global biogeochemical cycles (Bar-On and Milo 2019; Moran 2015). A fundamental paradox in marine ecology is to reconcile consistent ecosystem functions with the rapid and continuous turnover of microbial community composition (Louca et al. 2018). Ecosystem stability is traditionally attributed to functional redundancy under the assumption that multiple interchangeable taxonomic units perform identical metabolic tasks to provide a passive biological insurance against environmental fluctuations (Sunagawa et al. 2015; Allison and Martiny 2008). While this classical paradigm offers a theoretical buffer it increasingly fails to account for the highly structured and high frequency metabolic successions observed in dynamic coastal waters (Galand et al. 2018).

The perception of ecosystem stability remains deeply contingent on sampling resolution. Early annual and monthly time series focused on broad seasonal successions and initially reinforced the redundancy hypothesis by suggesting that shifting taxonomic groups fulfill interchangeable roles (Fuhrman et al. 2015; González-Motos et al. 2025). Subsequent diel scale studies sampling every three to six hours refined this perspective by revealing synchronized diurnal rhythms in autotrophs such as *Prochlorococcus* and photo-heterotrophs supported by daily phytoplankton exudates (Ottesen et al. 2013; Kieft et al. 2021). By artificially binning sequential specialists into persistent or co-occurring groups, low frequency observations are unable to unveil the discrete transitions underpinning ecosystem resilience (Ramond et al. 2025). Thus, more high frequency sampling is needed to fill this critical knowledge gap for better understanding of the coordinated temporal architecture of microbial functional networks.

Maintaining functional continuity amidst massive taxonomic turnover in dynamic coastal systems requires a mechanism more precise than passive redundancy, especially given the contributions of the rare biosphere (Sogin et al. 2006). To investigate this, we deployed an autonomous drone array technology in Daya Bay, China (Chen et al. 2024, 2026) to conduct metatranscriptomic sampling at a two-hour resolution over a seventy-two-hour period. Our results reveal that coastal stability arises not from static redundancy but from a deterministic and high-speed metabolic relay characterized by a radical restructuring of the active transcriptome where over 90% of gene clusters turn over every two hours.

This high frequency turnover drives a pattern we term taxonomic-functional locking where active community composition and specific metabolic repertoires exhibit an exceptionally strong coherency. Functional continuity is maintained through precise chronological successions of specialized organisms including a critical top-down orchestration by viral populations (Suttle 2007; Breitbart et al. 2018). Ultimately these findings demonstrate that coastal ecosystem resilience relies on active non redundancy where ecological functions are sequentially relayed by non-overlapping specialists optimized for distinct temporal micro-niches. This mechanistically deterministic framework fundamentally challenges the traditional species redundancy paradigm and redefines our understanding of microbiome stability in a changing ocean.

## Results

### Delineating the Temporal Metabolic Relay

To validate the paradigm of functional redundancy in coastal ecosystems, we investigated the transcriptional dynamics of the Daya Bay microbiome (**Fig. 1A**). Our high-resolution metatranscriptomic sampling (72 h at 2 h intervals; **Fig. 1B**) enabled the integration of protein-centric community-wide (CW) and genome-resolved (GR) (**Fig. 1C-D**) analytical strategies. While the CW approach encompassed all domains of life, the GR analysis targeted the main prokaryotic phyla and cyanophages. By capturing the intricate interplay between dominant prokaryotes and their viral predators, this dual strategy uncovers fine-scale metabolic shifts, thereby ensuring the robustness of the observed metabolic relay across distinct bioinformatic and taxonomic resolutions.

**Figure 1.**
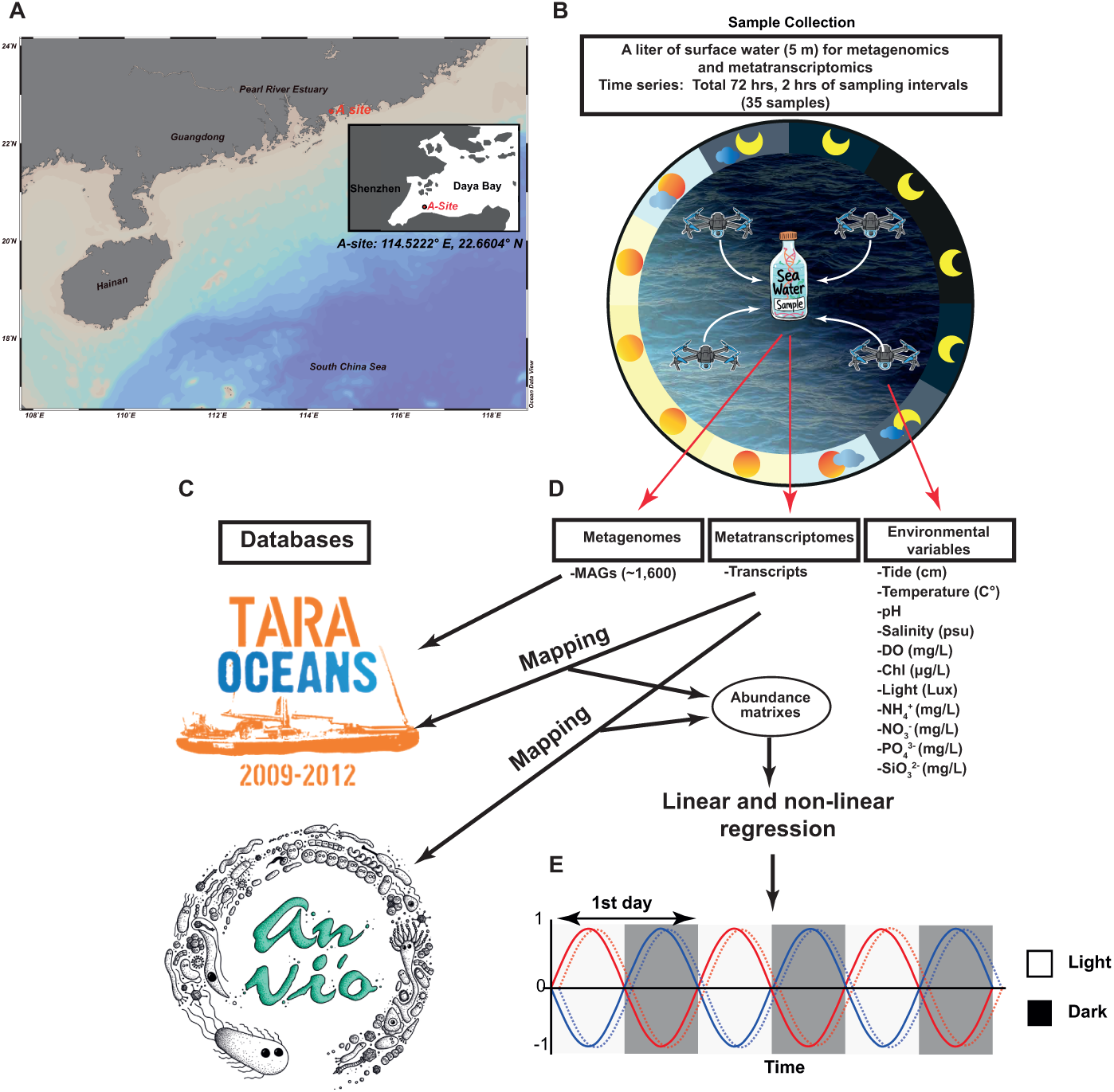
Overview of the experimental design and bioinformatics workflow. **A.** Geographic coordinates of Site A (114.5222° E, 22.6604° N) within Daya Bay, China. **B.** High-frequency autonomous sampling regime. Surface water (1-liter volume) was collected at a 5-m depth via an autonomous drone array, yielding 35 samples over a 72-hour period. The two-hour sampling interval captured three complete diurnal cycles. **C.** Dual analytical pipelines. Metatranscriptomic annotation utilized two independent databases to ensure cross-scale validation. **D.** Data integration framework. Metagenomic analysis leveraged approximately 1,600 Metagenome-Assembled Genomes (MAGs) for taxonomic anchoring, while metatranscriptomes enabled direct quantification of gene expression. Concurrent measurements of 13 physicochemical variables, including nutrients and irradiance, provided the environmental scaffold. Analytical modeling employed linear and nonlinear regressions to resolve temporal correlations from abundance matrices. **E.** Temporal modeling of diurnal oscillations. Conceptual sine waves illustrate the synchronization of microbial activities across light (white) and dark (grey) cycles over the 72-hour time series.

### Identification of Six Structured Transcriptional Modules

Multiple regression analysis (**Fig. 1E**) identified thousands of transcripts exhibiting significant and reproducible diurnal trajectories throughout the three-day cycle. These oscillations remained highly consistent by both approaches with 2,275 transcripts identified in the CW and 632 in the GR. Hierarchical clustering partitioned these dynamics into six discrete temporal modules per pipeline with Temporal Modules 1 to 6 for the CW approach and Temporal Modules A to F for the GR approach, respectively (**Fig. 2A-D**). Transcript counts ranged from 213 to 670 for the former approach and 37 to 211 for the latter approach (**Fig. S1**). The transcriptional weight of these modules, however, varied between these approaches: peak activity was identified during the light phase by the CW approach, whereas maximal specialization was identified during the nocturnal period by the GR approach (**Fig. S1**). Each module aligned with critical biogeochemical transitions such as the dawn to peak light shift or the evening organic scavenging phase, reflecting a strictly coordinated biological clock operating across the entire community (CW and GR).

**Fig 2.**
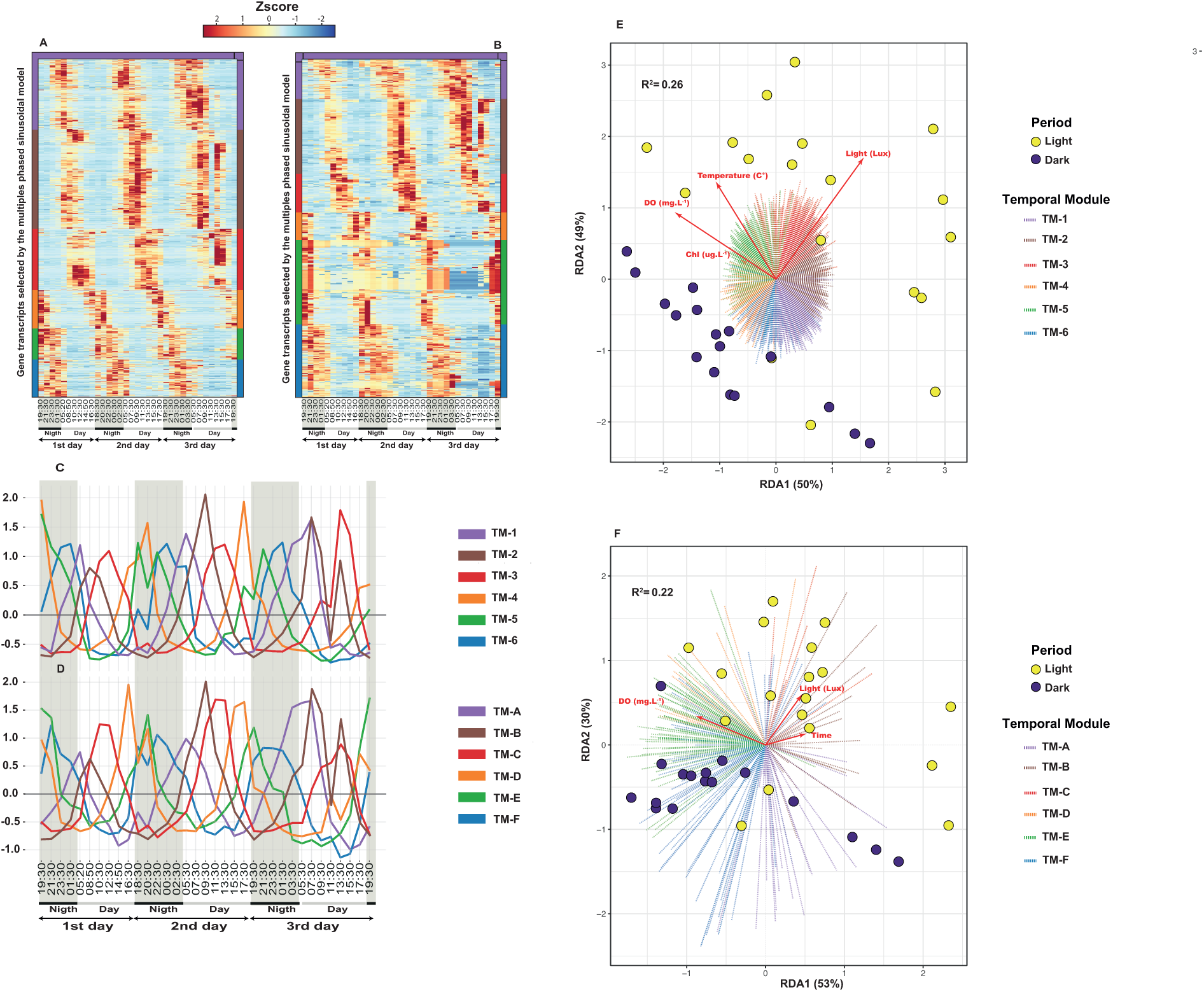
Synchronized diel rhythms with regular temporal offsets across a 3-day (72 hrs) period. Six distinct temporal modules (Temporal Modules 1 to 6 for the CW approach and Temporal Modules A to F for the GR approach, respectively) were identified using a clustering method with each module in relevance to a 2-3 h sampling interval. TM = Temporal Module. **A** and **B** are heatmaps of CW and GR approaches, respectively, with each line showing one gene transcript and each column the time of sampling. **C** and **D** are transcription profiles of Time Modules from the CW and. **E** and **F** are Redundancy analysis (RDA) between Temporal Modules of gene transcripts and environmental parameters, with each dot representing the sample dispersion and colored according to the light period based on Lux values. Red arrows indicate selected environmental parameters and the dashed lines a gene transcript colored according to the time cluster appurtenance.

### Sequential Turnover of the Diel Transcript Pool

The transition between these temporal niches involves a nearly complete replacement of the active transcript pool across the diel cycle. We categorized cycling transcripts into robust gene clusters using a sequence based clustering method via CD-HIT at 50% identity and 70% coverage. UpSet intersection analysis revealed a fundamental partitioning of this diel transcript pool, where 90% of active gene clusters are strictly restricted to a single Temporal Module with negligible overlap between successive phases (**Fig. S2**). This massive turnover is exemplified by the 181 and 118 unique clusters in Temporal Module 6 from the CW and GR approaches, respectively. Such results demonstrate that ecosystem stability is not sustained by a redundant background of constitutive expression but instead relies on the precise and non-overlapping transcription of specialized functional gene clusters. This high degree of phase exclusivity suggested an orchestrated temporal relay where each transcriptional assemblage was almost entirely replaced by a distinct specialist group to meet immediate environmental and metabolic constraints.

### Cross-Approach Validation and the Emergence of a Programmed Metabolic Continuum

To validate our temporal modules, we compared the CW and GR datasets using a cross-approach regression of Bray-Curtis dissimilarity matrices. The strong structural congruence confirmed that the observed successions are biologically driven, not artifacts of the bioinformatic pipeline. This comparison revealed strong structural congruence in all modules between the CW and GR approaches. For example, homologous time niches exhibited high coefficients of determination between community-wide Temporal Module 2 and genome-resolved Temporal Module B (*R*^2^ = 0.90) and between community wide Temporal Module 1 and genome resolved module Temporal Module A (*R*^2^ = 0.87) (**Fig. S3**). The resulting strong diagonal correlation across the comparison matrix proved that the chronological signal is an inherent biological property of the ecosystem that remained robust supported by both CW and GR approaches.

The overall architecture of the co-transcription network revealed an orchestrated temporal progression (**Fig. S4**). The network topology displayed a highly sequential and circular structure where transcripts spontaneously self-organized into an uninterrupted orbital trajectory over the 72-hour cycle. This circularity was underpinned by robust and statistically significant correlations with coefficients typically exceeding *R*^2^ > 0.70 for both Maximal Information Coefficient and linear or rank estimators (*p* < 10^-5^ after FDR correction) emerging exclusively between temporally adjacent clusters. Dawn-associated transcripts (Temporal Module 1 or Temporal Module A) occupy a central topological position, acting as an obligate bridge between the late-night assemblages and subsequent diurnal modules. The dense connectivity observed in the afternoon and nocturnal modules highlighted a programmed continuum where the transcriptional state of one metabolic niche acted as an obligate prerequisite for the next to ensure biogeochemical persistence through precise chronological handoffs of energy and nutrients.

### Deterministic Environmental Forcing of Circular Metabolic Successions

Redundancy Analysis (RDA) revealed that environmental cues, primarily light, temperature, dissolved oxygen, chlorophyll *a*, and time, significantly structured the successional modules, confirming the deterministic nature of these shifts. The selected environmental variables collectively explained 22–25% of the total transcriptional variance (*p* < 0.001), with light and dissolved oxygen emerging as the primary drivers within the model (**Fig. 2E-F**). The remaining variance may be governed by endogenous rhythms and biotic interactions, a hypothesis we further explore through the lens of viral-mediated metabolic shifts (see below).

Projected into the RDA space (**Fig. 2E-F**), transcriptional modules unfolded in an ordered succession, mapping directly onto environmental axes. The orthogonal orientation of light and dissolved oxygen vectors relative to late-day modules highlights a metabolic handoff, driven by the depletion of photosynthetic resources and the accumulation of byproducts. This chronological segregation in the RDA space provides robust evidence of strict temporal niche specialization, directly contradicting the paradigm of an active redundant metabolic landscape.

### Coordination between Taxonomic Identity and Diel Functional Potential

To identify the drivers of diel successions, we evaluated the congruence between taxonomic composition and diel functional profiles (**Fig. S5**). We observed a resolution-dependent coupling between community structure and functional repertoire, which challenges the active functional redundancy paradigm, the assumption that stochastic taxonomic turnover preserves global metabolic output (Galand et al. 2018).

Using the CW approach, significant coordination was observed at both taxonomic scales. At the class level taxonomic structure and functional profiles based on KEGG KOs were strongly correlated (Mantel *r* = 0.813, *p* < 0.0001) and accounted for 64% of the functional variance (*R*^2^ = 0.643). This coupling was strengthened at the genus level where community structure explained 66% of the functional distribution (*R*^2^ = 0.663), indicating that taxonomic identity remained a primary predictor of the diel gene pool even at the ecosystem-scale.

This deterministic signature became pronounced when focusing on dominant prokaryotic and cyanophage taxa (GR). While coordination at the class level showed high predictability (*R*^2^ = 0.745) comparison of functional and taxonomic profiles reached a near-linear congruence at the genus level where taxonomic identity predicted 96% of the functional response (*R*^2^ = 0.966, *p* < 0.0001; Mantel *r* = 0.981). Procrustes analysis further validated this symmetry (*M*^2^ = 0.012, *p* < 0.0001) as evidenced by the minimal residual vectors observed in the ordination space.

The consistent intensification of coupling from the class to the genus level demonstrates that microbial successions represent a high-precision relay where specific microbial populations strictly dictate the diel transcriptional program of the ecosystem. At the finest resolution the low *M*^2^ value reveals specific clade-bound metabolic coupling, which we define as taxonomic-functional locking. This provides direct mathematical evidence challenging the hypothesis of active functional redundancy. The sharp bipartite separation of samples highlighted a fundamental metabolic divergence between day and night, necessitating the independent treatment of diurnal and nocturnal functional profiles to resolve the biochemical signatures orchestrated by this microbial relay.

### Diurnal Metabolic Orchestration: A High-Precision Relay

The deterministic coupling revealed by Procrustes analysis suggests that Daya Bay successions represent a synchronized functional relay rather than stochastic taxonomic turnover. By integrating CW and GR signatures, we resolved the diurnal metabolic transitions (**Phases I–III; Fig. 3A**). Linking these transcripts to specific taxonomic entities and functional KOs (**Fig. 3B**) reveals that these transitions are not a turnover of redundant species, but a systematic progression from a stable metabolic backbone to highly specialized, non-overlapping gene deployments (**Fig. 3C**).

**Figure 3.**
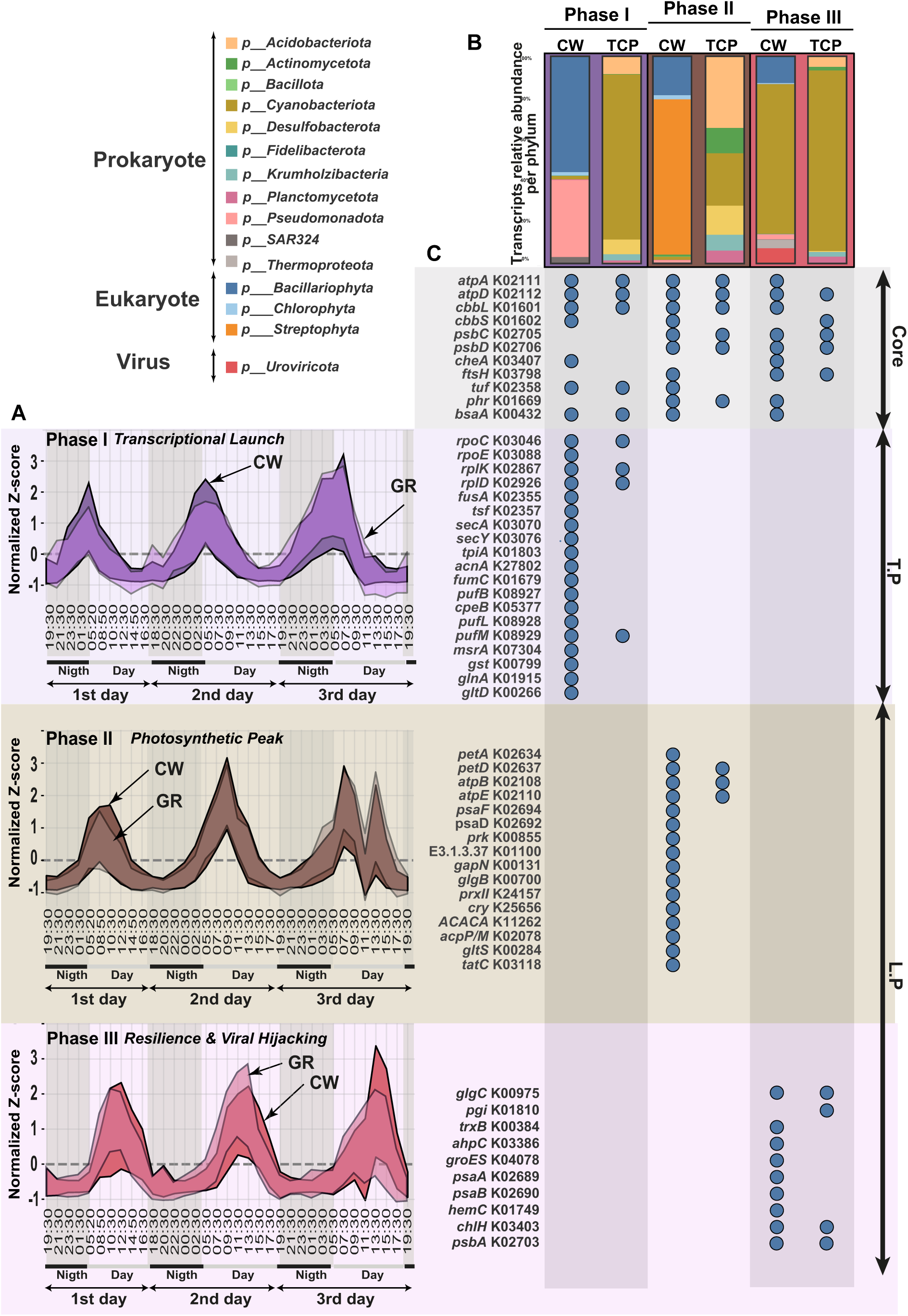
Taxonomic and functional specificity across transcriptional phases throughout the diurnal cycle. **A**. Expression profiles of temporal clusters. Curves represent *Z*-score normalized transcriptional dynamics over 72 hours. Overlapping pairs from both methodologies define specific phases, with CW profiles shown in darker shades and GR in lighter tones. Grey areas indicate nocturnal periods. **B**. Taxonomic composition of temporal clusters. Relative transcript abundance within prokaryotic, eukaryotic, and viral domains for each window. Bars compare community-wide (CW) and genome-resolved (GR) approaches, where color-coded phyla illustrate successions. **C.** Functional architecture and metabolic orchestration. Dot plot illustrating the presence of key functional genes across cycle phases. The metabolic core (grey) comprises genes expressed continuously, such as ATP synthase to ensure basal functions. The anticipation phase (purple) marks early activation of transcriptional and translational machinery during the dawn transition. The metabolic peak (brown) reflects the optimization of photosynthesis and energy conversion. Resilience and viral transition (red) involve the persistence of activity via stabilizing proteins and transcripts of viral origin to facilitate the shift to nocturnal conditions. L.P: light period; T.P: transition period.

While a constitutive functional core persisted throughout the light period, ensuring biogeochemical stability, it served primarily as a baseline infrastructure (**Fig. 3C, grey zone**). Energy production within this scaffold was maintained by the constant expression of ATP synthase alpha (K02111) and beta (K02112) subunits, while CO_2_ fixation potential remained anchored by Rubisco subunits (K01601, K01602). This shared background supported oxygenic photosynthesis via persistent transcription of Photosystem II core proteins (K02705, K02706) and magnesium chelatase (K03407), with cellular homeostasis further maintained by FtsH protease (K03798), elongation factor Tu (K02358), and essential defense genes including photolyase (K01669) and glutathione peroxidase (K00432). Beyond this shared functional core, the true engine of the biological carbon pump lay in phase-specific successions, with phase-exclusive functions accounting for 72%, 50%, and 46% of the total functional profile in Phases I, II, and III, respectively. This high degree of taxonomic-functional locking demonstrated that incoming taxa did not merely replicate existing efforts but occupied transient metabolic niches strictly synchronized with the solar progression.

### Phase I: The Transcriptional Launch

The dawn state was driven by a community-wide trophic mixing of early eukaryotes (54% of transcripts) and prokaryotes, primarily *Pseudomonadota* (32 %). From the genome-resolved approach, the dominance of *Cyanobacteriota* (54%) and *Bacteroidota* (31%) (**Fig. 3B**) completed an assemblage dedicated to the massive deployment of protein-manufacturing machinery. Robust transcription of RNA polymerase subunits (K03043, K03046) and ECF-family sigma factors (K03088) coincided with the synchronized expression of ribosomal genes (e.g., K02867, K02926), supported by elongation factors (K02355, K02357) and preprotein translocases (K03070, K03076). This rapid growth phase was fueled by the up-regulation of central carbon metabolism, including glycolysis (K01803) and the TCA cycle (K27802, K01679). Energy demands were met by a diverse photosynthetic apparatus, featuring light-harvesting complexes (K08927, K05377) and reaction centers (K08928, K08929), while oxidative stress protection (K07304, K00799) safeguarded the transition. Active nitrogen assimilation (K01915, K00266) provided the essential amino acid precursors required for eukaryotic biomass production (**Fig. 3C**).

### Phase II: The Photosynthetic Peak

Approaching maximum light intensity (11:00–12:00), the community shifted toward eukaryotic dominance (95% of activity), primarily driven by *Steptophyta* (72%) (CW) (**Fig. 3B**). This stage was optimized for maximal energy conversion and carbon sequestration, as evidenced by the high expression of the cytochrome *b6f* complex (K02634, K02637), F-type ATP synthase (K02108, K02110), and Photosystem I (K02694, K02692). Calvin cycle activity (K00855, K01100, K00131) was tightly coupled with polysaccharide storage (K00700) and thioredoxin-dependent photoprotection (K24157). Furthermore, cryptochrome detection (K25656) suggested a light-mediated fine-tuning of these metabolic shifts. Concurrently, the emergence of *Acidobacteriota* (33%, GR) aligned with transcriptomic signatures for fatty acid biosynthesis (K11262, K02078), amino acid metabolism (K00284), and the TatC translocase (K03118) (**Fig. 3C**).

### Phase III: Metabolic Momentum and Viral Hijacking

The late-morning transition was driven by the coupled dynamics of *Cyanobacteriota* (65%) and *Uroviricota* (6%) (**Fig. 3B**), functionally characterized by energy storage and stress response. Within this window, we detected a robust transcriptional pool of both bacterial and viral *psbA* (K02703) genes, marking the onset of targeted metabolic modulation. Enzymes such as glucose 1 phosphate adenylyltransferase (K00975) and glucose 6 phosphate isomerase (K01810) orchestrated the redirection of carbon flux toward carbohydrate accumulation. Simultaneously, a robust antioxidant suite was activated, including thioredoxin reductase (K00384), peroxiredoxin (K03386), and the GroES chaperonin (K04078). While photosynthetic transcription persisted through Photosystem I components (K02689, K02690) and porphyrin biosynthesis (K01749, K03403), this window was fundamentally distinguished by this viral modulation of the photosystem.

Structural discrimination of the transcriptomic dataset reveals tight metabolic relay dynamics between hosts and cyanophages. First derivative analysis (*d*Z/*d*x) of the *psbA* transcription pools demonstrates a precise kinetic synchrony between host and viral transcripts, challenging the standard model of passive viral replacement (**Fig. S6**). Rather than a staggered succession, the system exhibits synchronous metabolic momentum, where viral acceleration scales in real time with host velocity. This coupling peaks during a major transcriptional pulse (Samples 31–33), where the viral acceleration vector significantly outpaces that of the host, with viral acceleration reaching *d*Z/*d*x ∼5.5 and driving the viral transcript pool to surpass host expression levels. This proactive, hyper accelerated deployment indicates that cyanophages strategically synchronize and amplify their transcription to exploit maximal host energy fluxes, transforming these periods into a co-piloted transcriptional burst. The concurrent expression of an active viral encoded GIY YIG endonuclease confirms targeted host genome degradation and nucleotide recycling, while the confinement of viral capsid production to the subsequent nocturnal transition identifies this specific phase as a specialized, resource harvesting step prior to virion assembly (see next section).

### *The* Late Afternoon-Evening Functional Scaffold

As the community transitioned into the evening, the metabolic landscape shifted from energy acquisition to resource maintenance and priming. Analysis of the nocturnal temporal modules (**Phases IV–VI; Fig. 4A**) reveals that this stability is not a passive state, but a succession of specialized functional phases. Linking these transcriptional signatures to their taxonomic origins (**Fig. 4B**) confirms that nocturnal community dynamics are actively orchestrated. Differential transcript analysis identified a shared functional scaffold maintained throughout the night (**Fig. 4C**). This constitutive core ensured metabolic maintenance without light, anchored by the light-independent protochlorophyllide reductase system (K04037, K04038, K04039) and circadian period regulators (K26743). Energy maintenance within this framework integrated transhydrogenase (K00324) and glucose/mannose transporters (K17315) alongside constitutive DNA repair machinery such as *recA* (K03553) and *gyrB* (K02470). Beyond the second part of the day, Phases IV, V, and VI exhibited 61%, 64%, and 81% functional specificity, respectively.

**Figure 4.**
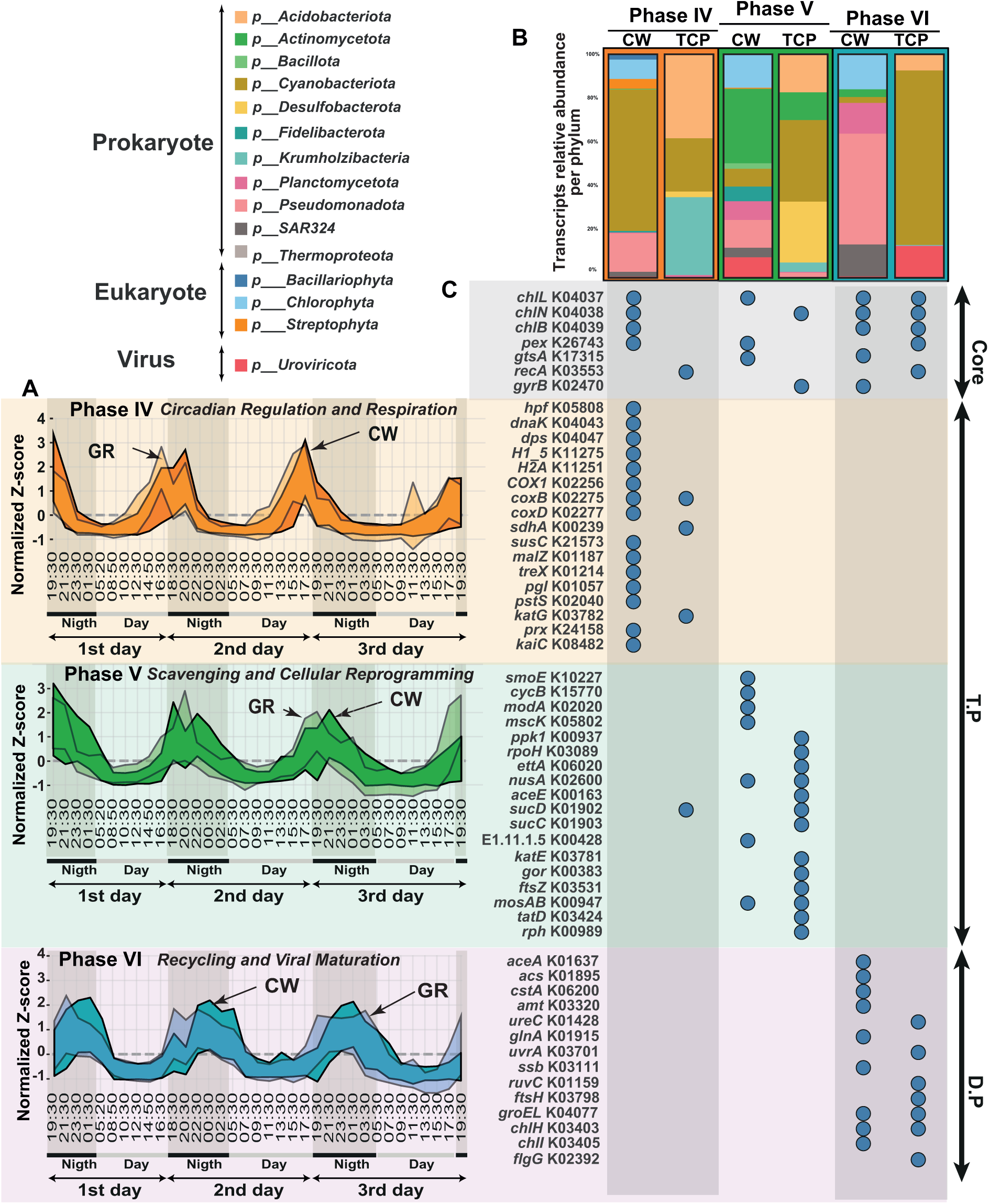
Taxonomic and functional specificity across the diurnal-nocturnal cycle. **A.** Expression profiles of temporal clusters. Normalized transcriptional dynamics (*Z*-score) over 72 hours for the six representative phases. Overlapping curves from both methodologies define specific temporal modules, with CW shown in darker shades and GR in lighter tones. Grey areas represent nocturnal periods. **B.** Taxonomic composition of temporal clusters. Relative transcript abundance within prokaryotic, eukaryotic, and viral domains. Bars compare CW approach and GR approach, with colors indicating major phyla to illustrate successions. **C.** Functional architecture and metabolic orchestration. Dot plot illustrating key KEGG Orthologs across cycle phases. The metabolic core (grey) comprises constitutively expressed genes such as Rubisco and ATP synthase. Respiratory and circadian priming (orange) involves circadian regulators and genome protection (e.g., *kaiC*, *dps*) for the dark transition. Scavenging and homeostasis (green) reflects membrane transporter diversification and energy reserve management. Nocturnal maintenance and recycling (pink) focuses on biomass recycling (e.g., glyoxylate cycle, urease) and viral structural assembly. T.P: transition period; D.P: dark period.

### Phase IV: Circadian Reprogramming and Respiratory Transition

The nocturnal phase initiated in the late afternoon, triggering a community-wide restructuring driven by *Cyanobacteriota* (52%, CW) and a tripartite GR contribution from *Acidobacteriota* (37%), *Cyanobacteriota* 24%), and *Krumholzibacteriota* (35%) (**Fig. 4B**). This pivot coincided with a sharp contraction of transcriptional entropy (Shannon Index H′), marking the shift toward tightened functional specialization (**Fig. S7A**). Induction of ribosome hibernation via *hpf* (K05808) signaled a deliberate throttling of protein synthesis. This slowdown was offset by a defensive deployment centered on the DnaK chaperone system (K04043) and genomic protection mechanisms (K04047, K11275, K11251). The energetic landscape concurrently abandoned anabolism for an activated respiratory chain, driven by cytochrome c oxidase subunits (K02256, K02275, K02277) and succinate dehydrogenase (K00239). Heterotrophic activity relied on carbon reserve mobilization via the starch-binding protein SusC (K21573) and various alpha-glucosidases (K01187, K01214, K01057). Meanwhile, phosphate transport (K02040) and a targeted antioxidant response (K03782, K24158) maintained redox homeostasis. This shift was functionally sealed by a rigorous circadian-metabolic coupling where KaiC oscillator expression (K08482) orchestrated the increasing respiration-to-translation ratio with mathematical precision (**Fig. S7B**).

### Phase V: Solute Scavenging and Cellular Reprogramming

The advanced transition phase, marked by the CW transcriptional dominance of *Actinomycetota* (32%), *Chlorophyta* (14%), and *Uroviricota* (9%), reflects a significant diversification of organic scavenging (**Fig. 4B**). This window is distinguished by the up-regulation of a broad-spectrum nocturnal “importome” featuring transporters for polyols (K10227), malto-oligosaccharides (K15770), molybdate (K02020), and potassium (K05802). This resource acquisition is coupled with the activation of polyphosphate kinase (K00937) for energy reserve management (**Fig. S8A**). Such metabolic synchronization is supported by a positive correlation between the integrated scavenging signal and energy storage transcripts (Spearman *ρ* = 0.535, *p* < 0.005; **Fig. S8B**), providing a mechanistic link between nocturnal uptake and the replenishment of community-wide energetic buffers.

Regulatory control during this period is evidenced by the expression of the stress-response sigma factor sigma32 (K03089), translation throttle proteins (*ettA*, K06020), and transcription termination factors (*nusA*, K02600). The enzymatic signature shifts toward central respiratory metabolism (K00163, K01902/03) and reinforced redox homeostasis (K00428, K03781, K00383). The community prepares for the subsequent growth cycle through the induction of the cell division protein FtsZ (K03531) and specialized molybdenum storage proteins (K00947). Active nucleic acid recycling via TatD DNase (K03424) and RNase PH (K00989) marks the completion of this cellular reprogramming sequence prior to the next dawn.

### Phase VI: Biomass Recycling and Viral Hijacking

Driven by a full-darkness community of *Pseudomonadota* (47%), *Chlorophyta* (15%), SAR324 (14) and *Planktomycetota* (13%) (CW), alongside a resurgence of *Cyanobacteriota* (74%) (GR) (**Fig. 4B**), this phase transitioned toward systemic biomass recycling. This shift was mediated by the glyoxylate cycle (isocitrate lyase, K01637) and acetyl-CoA synthesis (K01895), with the induction of carbon starvation proteins (K06200) ensuring survival via carbon recovery from endogenous reserves.

Heterotrophic activity was coupled with a robust nitrogen acquisition signature including ammonium transporters (K03320), urease (K01428), and glutamine synthetase (K01915). Cellular maintenance was supported by high expression of excision repair (K03701), strand protection (K03111, K01159), and the FtsH-GroEL system (K03798, K04077). Sustained transcription of pigment biosynthesis genes (K03403, K03405) and flagellar motility machinery (K02392) suggested strategic preparation for dawn.

A highly structured co-expression network further defined this transition (**Fig. S9**). The integration of viral capsid production with host nitrogen metabolism was evidenced by positive synchronization between capsid proteins, glutamine synthetase (*glnA*), and ammonium transporters (*amt*). In contrast, the dense negative correlation network revealed a viral-induced redirection of cellular resources, where capsid maturation served as a central hub antagonistic to host energetic maintenance, showing significant anti-correlation with ATP synthase alpha (*atpA*) and NADH dehydrogenase (*ndh*).

### Structural and Phylogenetic Validation of the *psbA* Pool

We resolved the taxonomic origin of the diel *psbA* pool (K02703) using a multi-scale structural validation pipeline based on the ESM-2 protein language model. After benchmarking various model architectures (**Fig. S10**), we established that prediction consistency and accuracy plateaued with the 650M parameter model. Using a weighted consensus adjudicator that integrates the 35M, 150M, and 650M expert models, we partitioned 233 candidate sequences into distinct evolutionary cohorts. Our analysis identified 33.7% of the sequences as viral and 53.8% as non-viral, while 12.5% were classified within a “Grey Zone”(likely representing rapidly evolving viral clades) and 7.7% of low-confidence sequences were excluded (**Fig. S11**). Phylogenetic reconstruction (**Fig. S12**) confirms this architecture, as viral sequences, including those identified in the “Grey Zone,” nest robustly within established cyanophage clades and form a monophyletic group distinct from the cyanobacterial outgroup. This convergence between structural signatures and phylogenetic signal validates the robustness of our annotation. Our results demonstrate that the apparent redundancy of the *psbA* pool masks a complex taxonomic partitioning, where cyanophages deploy structurally optimized variants to meet specific metabolic requirements during infection, rather than merely supplementing host functions.

## Discussion

Marine ecosystem stability is traditionally pinned on functional redundancy, a framework where high taxonomic diversity operates as a biological insurance policy (Yachi and Loreau 1999; Louca et al. 2018). Our findings subvert this assumption. High-frequency metatranscriptomic tracking reveals a volatile reality where up to 90% of active gene cluster identities are replaced at each phase transition, shifting from energy acquisition in the light to resource maintenance and priming in the dark. This is not a passive turnover but a deterministic, six-phase relay (Phases I–VI) that provides the fundamental insurance against environmental variability.

The daily cycle begins with Dawn Metabolic Activation in Phase I or Module 1 (**M1, Fig. 5A**), a structural reset where the induction of the transcriptional apparatus (K03043, K03088) and ribosomal biogenesis establish readiness. This leads to the Calvin Cycle-Driven Carbon Surge in Phase II (**M2**), a peak-light commitment to photosynthetic throughput (K02634, K02637, K02694) and organic biomass synthesis. True stochastic redundancy would naturally decouple community structure from this diel metabolic pool (Galand et al. 2018), yet our Procrustes analysis demonstrates a near-linear coordination between taxonomic and functional profiles (*R*^2^ = 0.966). This internal clockwork is actively governed by the circadian regulator KaiC (K08482), which prioritizes maintenance over translation (Spearman *ρ* = 0.828, *p* < 0.001; **Fig. S7B**), rather than by passive environmental forcing.

**Figure 5.**
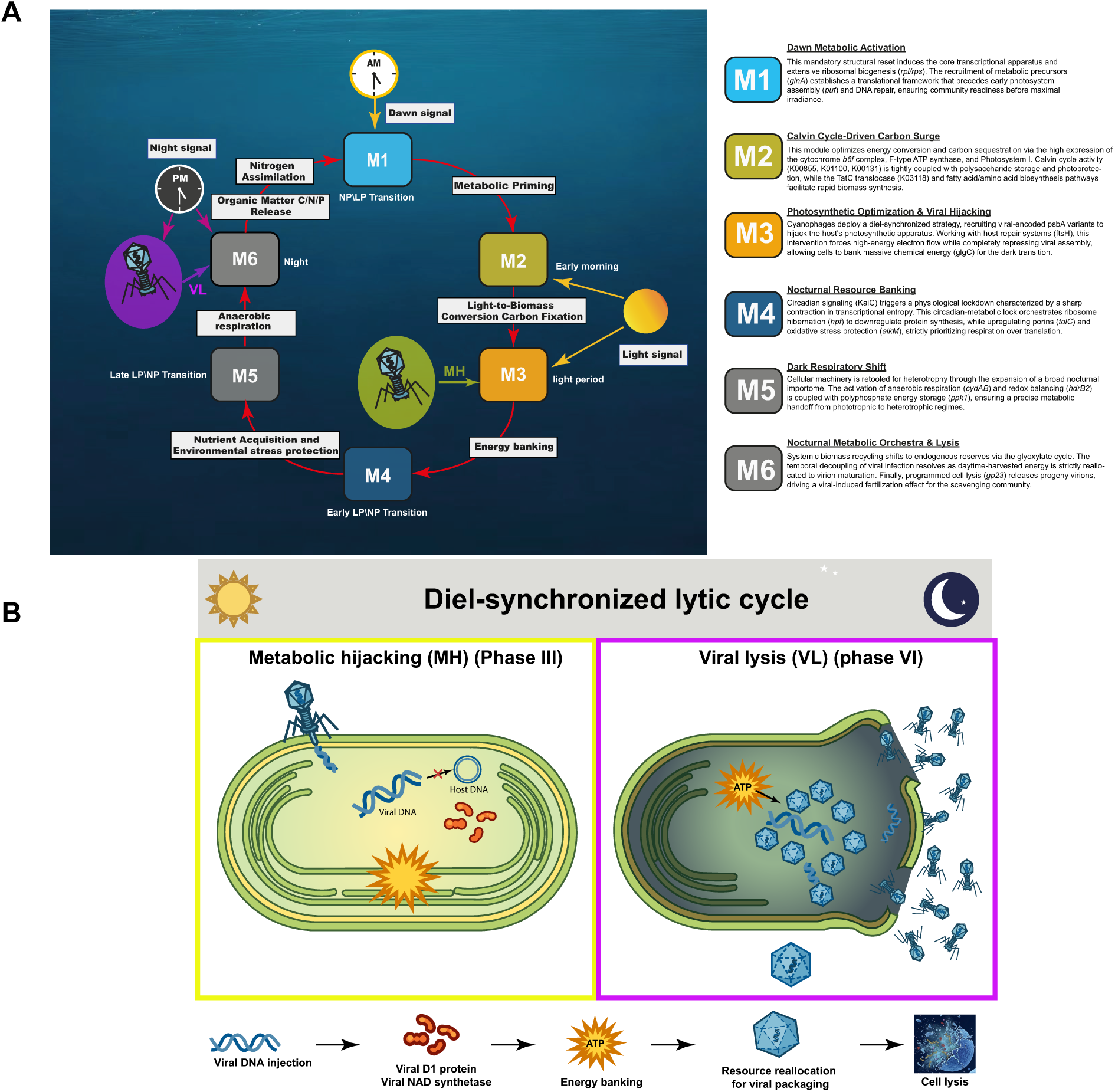
Integrated framework of the diel metabolic cascade and the cyanophage lytic cycle. **A**. Chronometric relay of the six functional modules (M1–M6) governing the coastal diel metabolic cycle. Solid black arrows indicate the primary chronological sequence; the dashed arrow highlights the ecological feedback loop where viral-induced cell lysis at the end of the dark period (M6) releases labile organic matter (C/N/P) to fertilize the system, effectively closing the ecosystem loop. The causal activation sequence (M1 → M2 → M3 → M4 → M5 → M6) represents a highly conserved functional relay strictly synchronized with diurnal solar irradiance and nocturnal biophysical constraints. **B.** Conceptual model of the diel-synchronized lytic cycle orchestrated by cyanophages. (Left) Daytime Metabolic Hijacking in Phase III: Upon infection, cyanophages inject their DNA but deliberately delay structural assembly. Instead, the viral genome remains episomal, excluding a dormant lysogenic state and actively transcribes auxiliary metabolic genes (AMGs), including *psbA* (encoding the viral D1 protein). These viral proteins target the host’s thylakoid membranes to sustain photosynthetic momentum and maximize intracellular energy accumulation (glycogen banking) during peak irradiance. (Right) Nocturnal Assembly and Lysis in Phase VI: As the ecosystem transitions into darkness, the accumulated energetic resources are systematically reallocated toward viral structural biosynthesis. Capsid maturation and DNA packaging culminate in programmed host cell lysis (*gp23*), releasing progeny virions prior to the subsequent dawn. This temporal decoupling maximizes the photosynthetic yield of the host before completing the lytic cycle.

The microbial successions act as a high-precision relay optimized for energy flux, following an invariant blueprint dictated by thermodynamic laws (Kleidon 2023). A contraction of transcriptional entropy (H’) marks the transition into the nocturnal phases (**Fig. S7A**), where the community compresses its repertoire into a specialized survival program. During Nocturnal Resource Banking in Phase IV (**M4**) and the Dark Respiratory Shift in Phase V (**M5**), the community retools for heterotrophy via the importome (K10227) and polyphosphate energy storage (K00937, *ppk1*). This sequential dependency, where each phase establishes the chemical prerequisite for the next (Braakman et al. 2017), bypasses resource competition while maximizing photon yield (Muratore et al. 2022). This self-organized resilience challenges conventional ecological modeling (González-Motos et al. 2025).

Our results fundamentally recast marine phages as active facilitators of this relay. During Photosynthetic Optimization and Viral Hijacking in Phase III (**M3**), cyanophages deploy structurally distinct, deep learning-adjudicated *psbA* variants (K02703) to hijack the host photosynthetic apparatus (Sullivan et al. 2006; Kieft et al. 2021). First-derivative analysis (*d*Z/*d*x) of the environmental *psbA* pool provides mathematical proof of this kinetic coupling, as viral transcript acceleration scales in real time with host velocity, peaking at irradiance (*d*Z/*d*x ∼ 6; **Fig. S6**). This daylight hijacking forces massive glycogen banking (*glgC*), an energy capital that the virus only calls upon during the nocturnal transition. The cycle culminates in the Metabolic Orchestra & Lysis in Phase VI (**M6**), where the temporal decoupling occurs: chemical energy harvested in Phase III is reallocated toward virion maturation and gp23-mediated capsid assembly (**Fig. 5B**). Finally, programmed cell lysis provides a viral-induced fertilization effect, spraying labile organic matter into the water column to support the nocturnal importome (**Fig. 5A**).

Moving from broad metabolic potential to acute taxonomic-functional locking can change how we model microbial procssess in the ocean. If every temporal niche is coupled to a specific biological actor, the biological carbon pump may be far more sensitive to community desynchronization than previously known (Allison and Martiny 2008). The threat of climate disruption is not just about species extinction, but the desynchronization of the global metabolic clockwork—a silent uncoupling now quantifiable through transcriptional entropy, circadian locking, and viral kinetic derivatives.

While this study provides a robust high-resolution view of coastal dynamics, the observed metabolic relay may operate differently in nutrient-poor open ocean environments, where viral influence and resource competition dynamics could differ. Future studies should extend this temporal tracking across distinct biogeochemical provinces to determine the universality of this deterministic clockwork. By identifying these specific temporal bottlenecks, we provide a new diagnostic framework for ocean forecasting, shifting the focus from mere presence-absence diversity to the preservation of precise, time-locked functional orchestration.

## Materials and Methods

### High-Frequency Sampling and Environmental Context

Coastal surface seawater samples (n = 35) were collected from Station A (22.6604328°N, 114.5221549°E) in Daya Bay, South China Sea, over 72 consecutive hours (October 28–31, 2021). Sampling was executed at two-hour intervals using an autonomous drone array (Egretta QC EMC50 Pro) as detailed by Chen et al. (2026). Hydrographic parameters (temperature, light, dissolved oxygen, chlorophyll-*a*) were recorded concurrently via drone-mounted sensors. Seawater was filtered through 0.2 µm membranes for cellular fractions and 0.02 µm membranes for viral fractions, followed by immediate flash-freezing in liquid nitrogen.

### Metatranscriptomic Library Preparation

RNA was extracted from 0.2 µm membranes using the FastDNA SPIN Kit (MP Biomedical), followed by DNase I treatment (TaKaRa) and purification via the RNeasy MinElute system (QIAGEN). Although SortMeRNA v4.3.6 was used for initial ribosomal RNA depletion, the complete transcriptional pool was retained to capture active ribosomal biogenesis and core translational machinery dynamics. cDNA libraries were sequenced on an Illumina NovaSeq 6000 (2×150 bp), yielding 4.7 TB of raw transcriptomic data.

### Community-Wide Protein-Centric Approach

To circumvent assembly biases, we constructed a reference protein catalog integrating the TARA Oceans gene atlas, 700 Daya Bay metagenome-assembled genomes (MAGs) (Chen et al., 2026), and targeted South China Sea MAGs (Xu et al. 2024). The dataset was clustered at 90% identity using CD-HIT (Li et Godzik 2006). Prokaryotic taxonomy was resolved via GTDB-Tk v2.4.0 (Chaumeil et al. 2020) and DIAMOND (Buchfink et al. 2015) against a curated 140,000-genome database and nr from NCBI. Viral functions were resolved using DeepKOALA (Yu et al. 2026), KofamKOALA (Aramaki et al. 2020) and InterProScan (Quevillon et al. 2005). Reads were mapped against the catalog via DIAMOND, requiring a 90% alignment identity and coverage threshold.

### Genome-Resolved Approach and Pangenome Mapping

Population-level activity was tracked using a pangenome framework (Ottesen et al. 2013) within anvi’o (Eren et al. 2015). For 13 distinct taxonomic clusters (Table S1), pangenomes were curated and annotated via DeepKOALA, KofamKOALA and InterProScan. Metatranscriptomic reads were aligned against pangenome references using DIAMOND blastx mode, retaining alignments with ≥70% sequence identity and E-value <10⁻⁵. Counts were normalized to relative abundance for cross-sample comparison.

### Identification of Diurnal Cycling Genes and Modular Clustering

Transcriptional oscillations were identified by screening against a library of 60 synthetic sinusoidal models (10-minute sliding phase offsets) using mictools (Albanese et al. 2018). We required a Maximal Information Coefficient (MIC) >0.69 (Reshef et al. 2011) and a Pearson/Spearman correlation >0.69 (FDR-corrected p<0.05). Significantly cycling transcripts were *Z*-score transformed and grouped into six chronological modules (Phases I–VI) using complete-linkage hierarchical clustering based on correlation distances.

### Taxonomic-Functional Locking and Structural Adjudication

Congruence between community structure and metabolic potential was validated via Mantel tests and Procrustes analysis (PROTEST) (Peres-Neto et Jackson 2001); using Bray-Curtis dissimilarity matrices across taxonomic ranks. For *psbA* (K02703) variants, we developed a structural validation pipeline using ESM-2 protein language models (Lin et al. 2023). Unlike sequence-homology-based methods, ESM-2 captures structural motifs indicative of evolutionary descent. We fine-tuned architectures (8 M to 650 M parameters) using a ground-truth dataset of verified viral and cyanobacterial sequences. The 650 M architecture was established as the final adjudicator (**Fig. S11**) based on superior precision-recall metrics. Ambiguous “Grey Zone” sequences were segregated from high-confidence assignments (softmax score ≥0.85) and validated via maximum-likelihood phylogenetic inference (IQ-TREE v2.0.3; LG+R10 model).

### Statistical and Computational Framework

Transcriptional complexity was assessed using Shannon entropy (H’). Circadian-metabolic coupling was evaluated by correlating CLR-transformed expression of *kaiC* (K08482) against a metabolic differential representing the ratio between respiratory and translational machinery. All statistical analyses were performed using custom Python pipelines (Pandas/SciPy), with CLR transformation applied to eliminate compositional interdependence biases.

## Acknowlegements

This study was supported by the National Natural Science Foundation of China (Nos. W2533069, 42321004), the Ocean Negative Carbon Emissions (ONCE) program, the Shenzhen Key Laboratory of Marine Archaea Geo-Omics, Southern University of Science and Technology (SUSTech) (SYSPG20241211173725010), Shenzhen Science and Technology Innovation Program (Grant No. JCYJ20220530115401003). Computation in this study was supported by the SUSTech Center for Computational Science and Engineering. This paper contributes to the Science Plan of the UN Ocean Decade Global Ocean Negative Carbon Emissions (Global ONCE) Program, its first project iCUBEs (#52.2), and Shenzhen Ocean University, Shenzhen 518055, China.

