## Supplemental figures for "Active non-redundancy and viral orchestration sustain diel microbial successions in the coastal ocean"

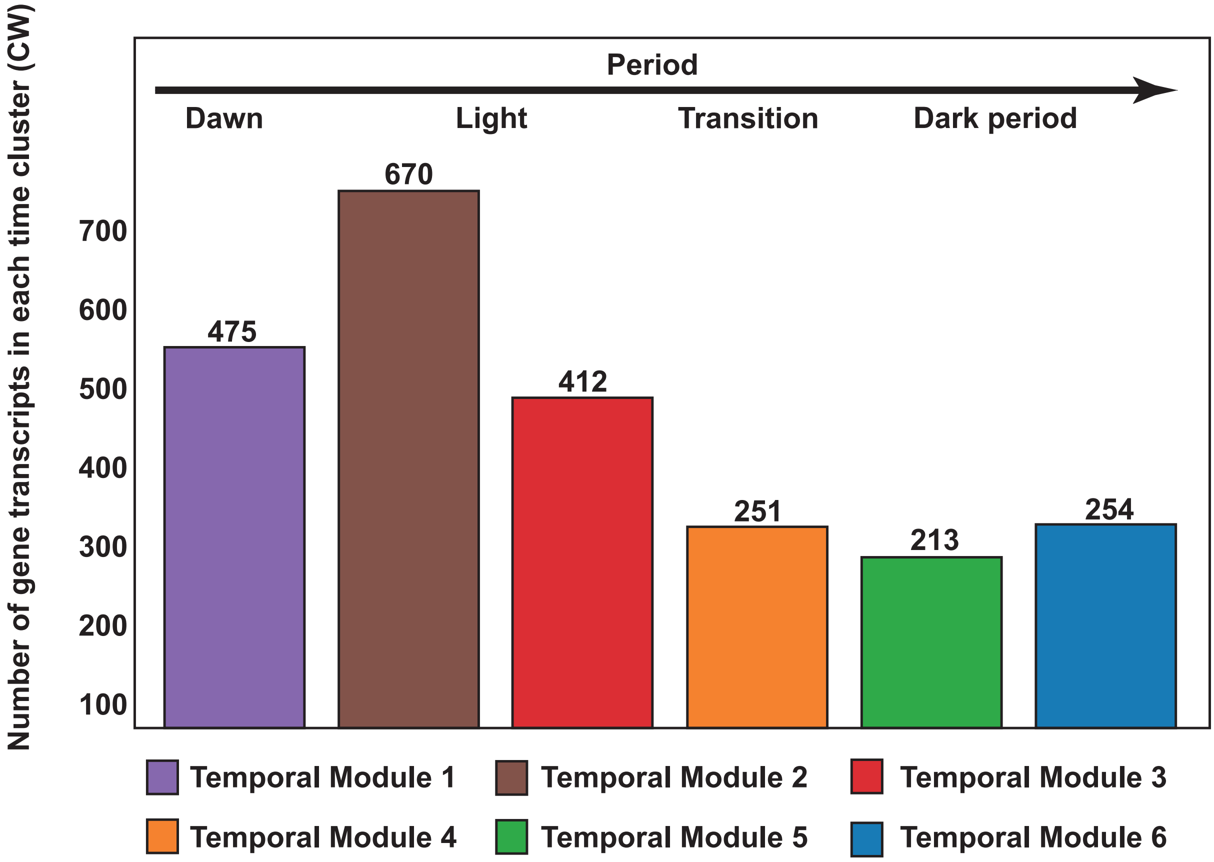
Number of gene transcripts in each time cluster (CW)
Period
Dawn
Light
Transition
Dark period
700
600
500
400
300
200
100
475
670
412
251
213
254
Temporal Module 1
Temporal Module 2
Temporal Module 3
Temporal Module 4
Temporal Module 5
Temporal Module 6

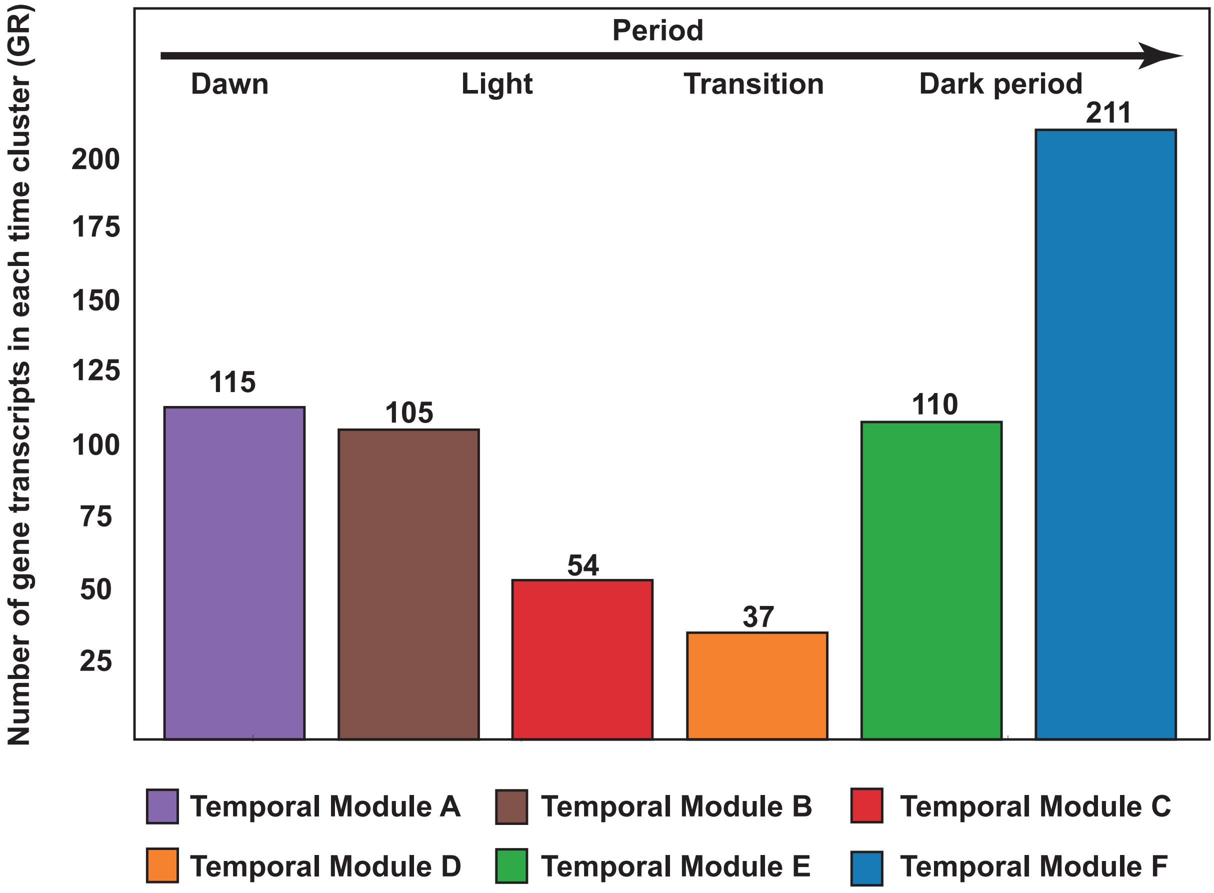
Number of gene transcripts in each time cluster (GR)
Period
Dawn
Light
Transition
Dark period
200
175
150
125
100
75
50
25
115
105
54
37
110
211
Temporal Module A
Temporal Module B
Temporal Module C
Temporal Module D
Temporal Module E
Temporal Module F

1

**Figure S1 | Magnitude and distribution of the active transcriptional repertoire across Temporal Modules. CW = Community-Wide; GR = Genome-Resolved.**

**A–B.** Quantification of unique gene transcripts within chronologically ordered functional modules for the community-wide (top) and genome-resolved (bottom) approaches. Transcript counts varied from 213 to 670 in the community-wide approach and 37 to 211 within the genome-resolved approach, illustrating the distinct metabolic weight of each temporal niche. Peak activity in the CW approach aligned with the day light period (Temporal Module 2, 670 gene transcripts) whereas the GR approach identifies maximal specialization during the nocturnal period (Temporal Module F, 211 gene transcripts). This asymmetry necessitated the integration of both methodologies to fully resolve the complexity of the coastal metabolic relay.

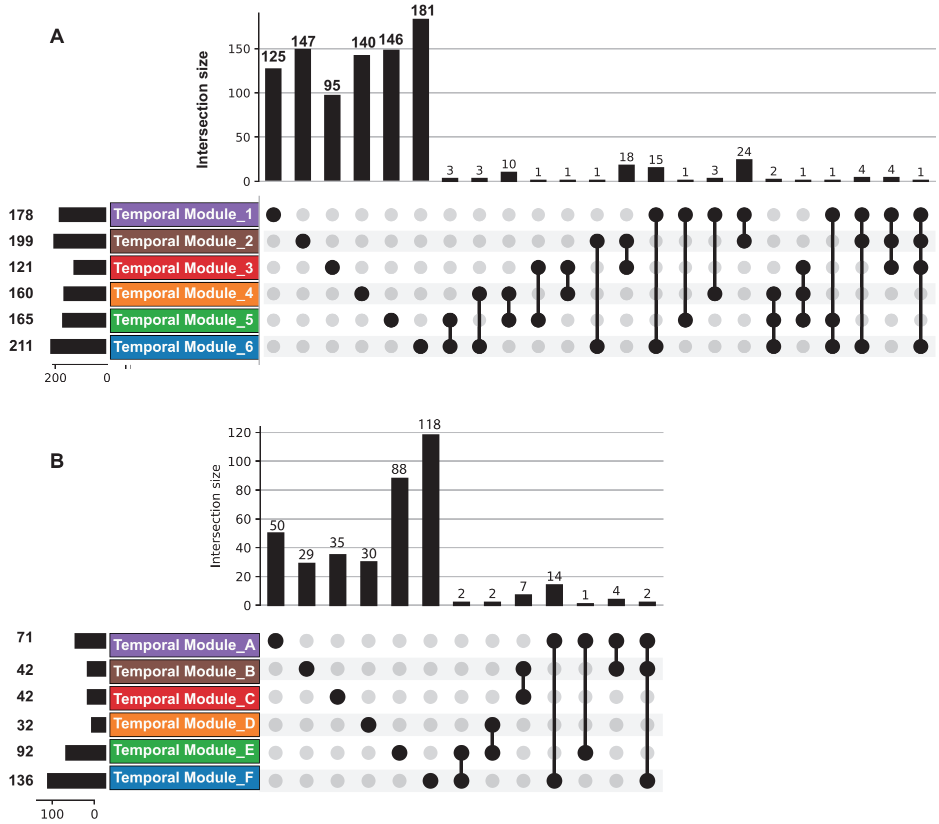


**Figure S2 | High frequency turnover of active gene clusters across Temporal Modules.**

**A–B.** UpSet plot analysis illustrating the intersection and unique distribution of cycling gene clusters across six Temporal Modules for the community wide (**A**) and genome resolved (**B**) approaches. In both approaches transcriptional activity was defined by extreme phase specificity where approximately 90% of active clusters remained strictly restricted to a single temporal module. The dominance of these single set columns compared to the negligible multi module intersections confirmed a massive turnover and minimal functional overlap between successive diel phases. While the total number of clusters varied between the community-wide and genome resolved approaches, the convergence of these independent methodologies toward a radical partitioning of the diel transcript pool provided robust evidence of an orchestrated metabolic relay. Within this framework specialized modules are exclusively activated to match the immediate environmental and biogeochemical constraints of the Daya Bay.

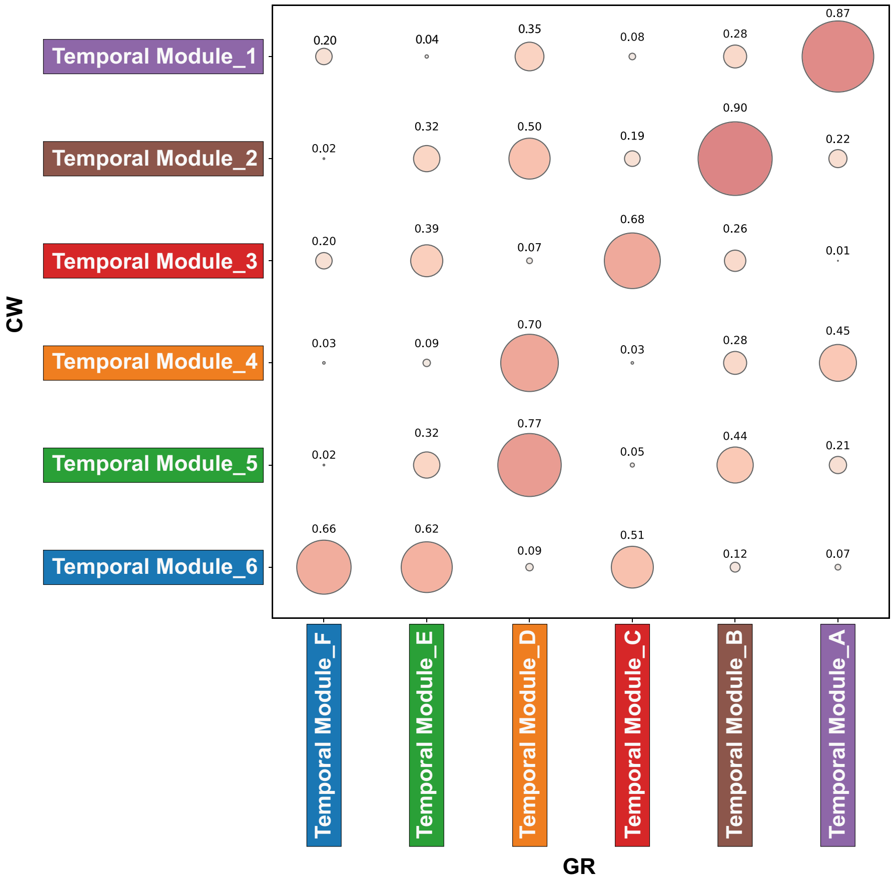


**Figure S3 | Structural congruence matrix between community wide and genome resolved approaches.**

Correlogram illustrating the statistical coordination between community-wide (y axis) and genome-resolved (x axis) Temporal Modules. The matrix is based on the correlation of Bray Curtis distance profiles calculated from CLR normalized transcript counts. Each cell displayed the coefficient of determination ($R^2$) where the size and color intensity of the circles represented the strength of the linear relationship between the two methodologies. Homologous modules exhibited a strong diagonal correlation which validated the robustness of the chronological signal across both protein centric from community-wide and genome-resolved scales.

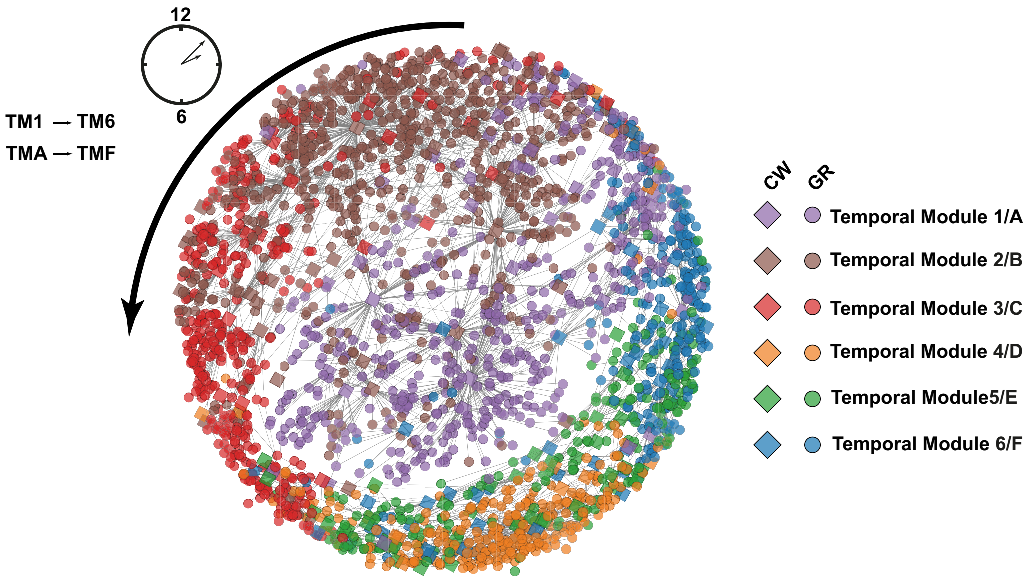


**Figure S4 | Orbital architecture and sequential trajectory of the co-transcription network.**

Network reconstruction using an edge weighted spring embedded layout based on the Maximal Information Coefficient (MIC) to visualize linear and non-linear dependencies within the complete community. Within this architectural framework diamonds represent transcripts from the genome-resolved (GR) dataset while circles denote the community-wide (CW) dataset. Node colors correspond to the six discrete Temporal Modules identified across the 72-hour cycle ranging from TM-1/A to 6/F. A symbolic clock and directional arc illustrate the chronological progression and the strictly sequential nature of the orbital trajectory. The resulting circular topology is sustained by robust and statistically significant co-transcription correlations with coefficients typically exceeding $R^2 > 0.70$ for MIC and Pearson or MIC and Spearman estimators ($p < 10^{-5}$ after FDR correction) occurring predominantly between temporally adjacent clusters. Purple nodes representing dawn associated transcripts (Temporal Module 1/A) act as a topological bridge to facilitate the transition toward diurnal modules and subsequent progression through the afternoon and nocturnal phases (TM-4/D, 5/E, and 6/F).

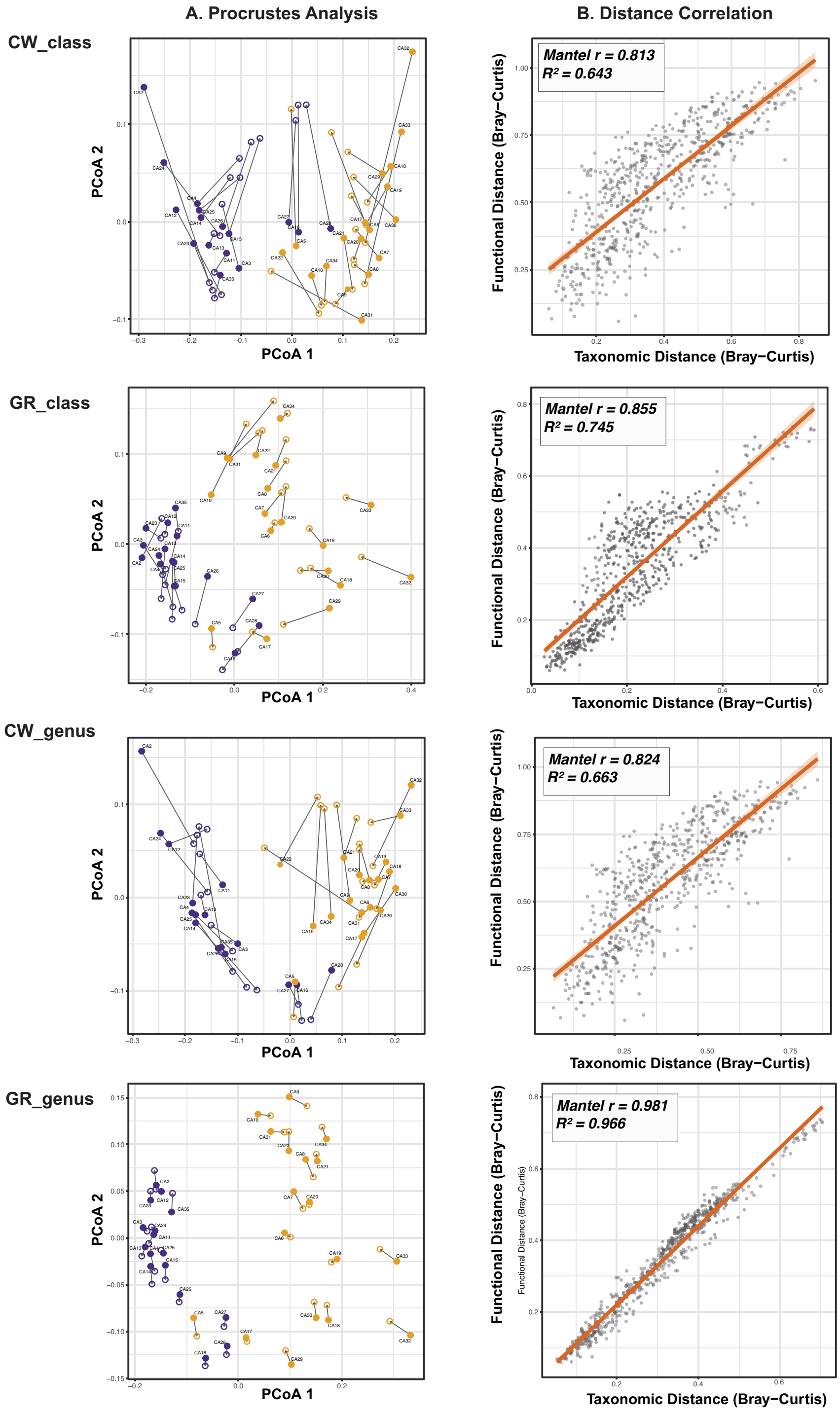


**Figure S5 | Multi-scale convergence of taxonomic structure and functional potential.**

**A.** Procrustes analysis ordinations illustrating structural symmetry between taxonomic and functional distance matrices across scales. The analysis compares the community-wide and genome-resolved approaches at both class and genus resolutions. Short residual vectors and clear bipartite separation across all panels confirmed fundamentally distinct and highly coordinated metabolic regimes between light (yellow) and dark (purple) periods. **B.** Distance correlation plots showing linear regressions of flattened Bray-Curtis distance matrices. The coupling between taxonomic identity and active functional profiles intensified systematically with increasing resolution, transitioning from a broad correlation at the community level to a near-linear deterministic relationship at the genus scale. This progressive tightening of the regression along the diagonal provided a robust mathematical challenge to the functional redundancy hypothesis, demonstrating that taxonomic identity served as a primary predictor of the ecosystem’s transcriptional program.

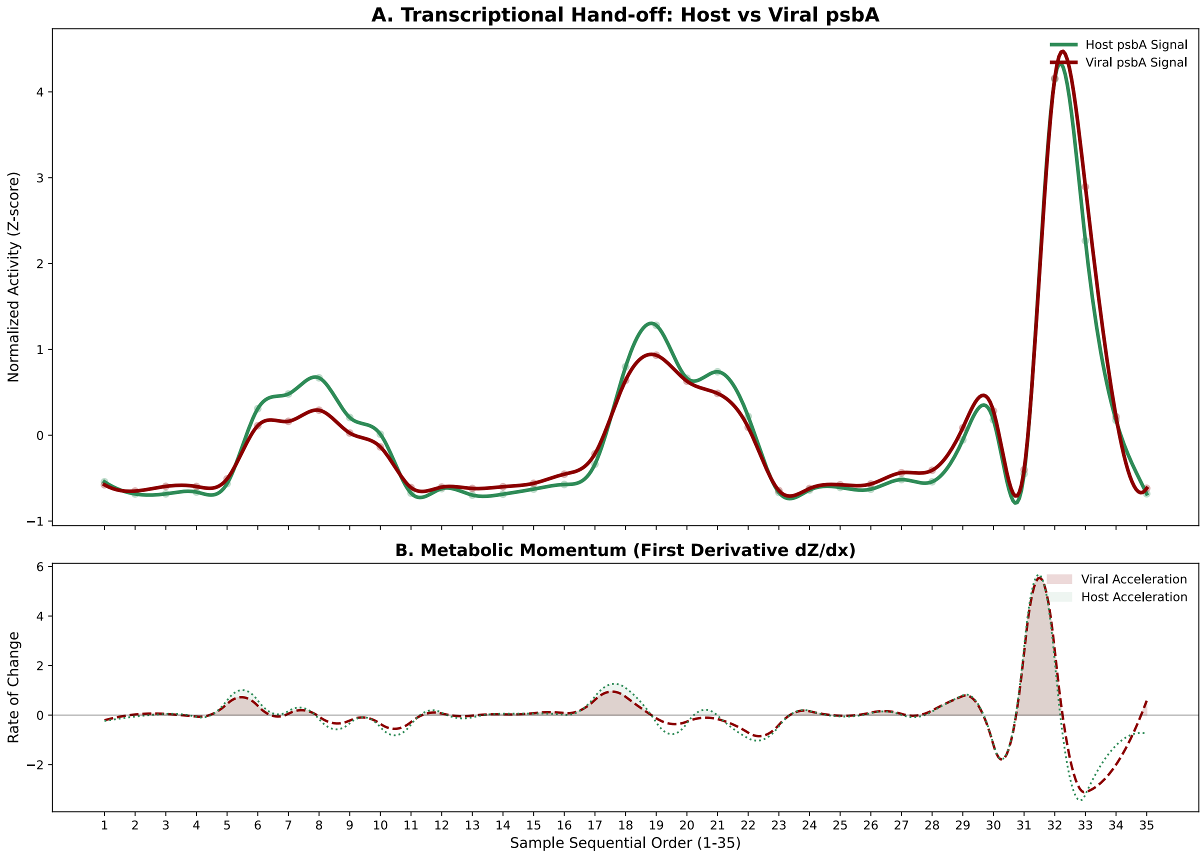


**Figure S6 | Kinetic analysis and momentum of the single-copy reaction center gene *psbA*.** **A.** Relative activity (Z-score normalized transcript counts) of the *psbA* pool from *Cyanobacteriota* (Host) and *Uroviricota* (Virus) across the high-resolution temporal modules. While absolute abundance profiles (left panels) suggest a passive succession, normalized activity (Z-scores) reveals a robust near-perfect synchrony between host repair and viral deployment. **B.** First-derivative analysis (dZ/dx) resolves the metabolic momentum (rate of change) of the *psbA* pool. Far from a delayed replacement, viral acceleration scales in real-time with host velocity, spiking massively ($dZ/dx \sim 5.5$) at peak irradiance (Sample 31; day 3, 11:30) when the host endogenous repair machinery saturates. This proactive synchronization implies a diel-synchronized viral hijacking to maintain photosynthetic throughput under maximal energy fluxes.

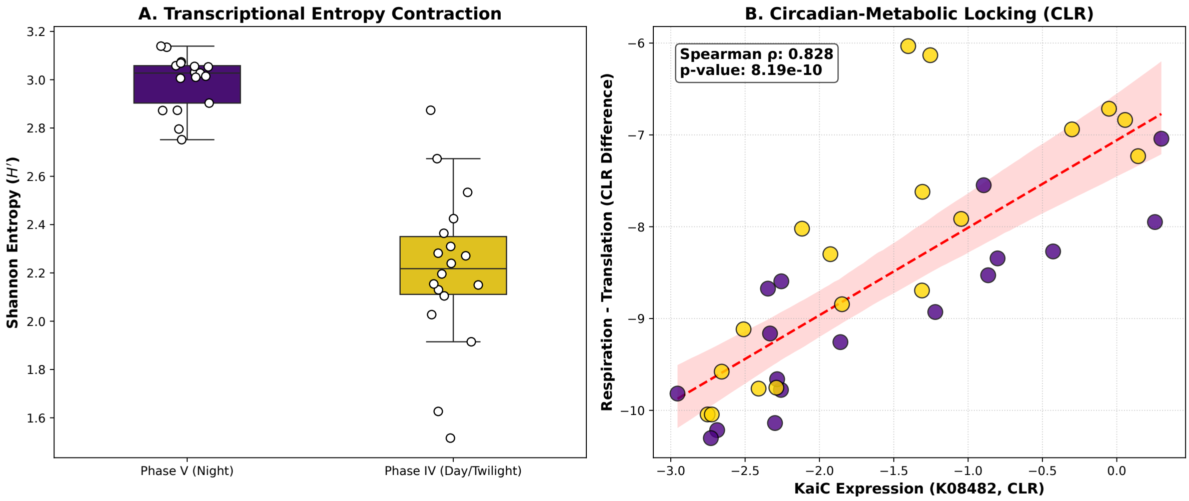


**Figure S7 | Informational and mechanistic coupling of the nocturnal metabolic transition.**

**A.** Transcriptional entropy contraction. Boxplot and individual data points showing the Shannon entropy (H') of the community wide transcriptomic profile across phases. The significant reduction in entropy during the light to dark transition (yellow) compared to the nocturnal state (purple) quantifies a deterministic functional tightening. This shift reflects a move from expansive metabolic diversification toward a specialized survival program centered on genome protection and protein maintenance. **B.** Circadian metabolic coupling. Scatter plot illustrating the high-fidelity coordination between the expression of the master circadian regulator KaiC (K08482) and the metabolic ratio of respiration (cytochrome c oxidase) to translation (ribosomal proteins). The robust positive correlation (Spearman *rho* = 0.859, $p < 0.0001$) demonstrates that the strategic downregulation of protein synthesis in favor of respiratory maintenance is governed by the biological clock. Yellow points representing Day/Twilight and purple points representing Night reveal that metabolic locking initiates during the late afternoon transition.

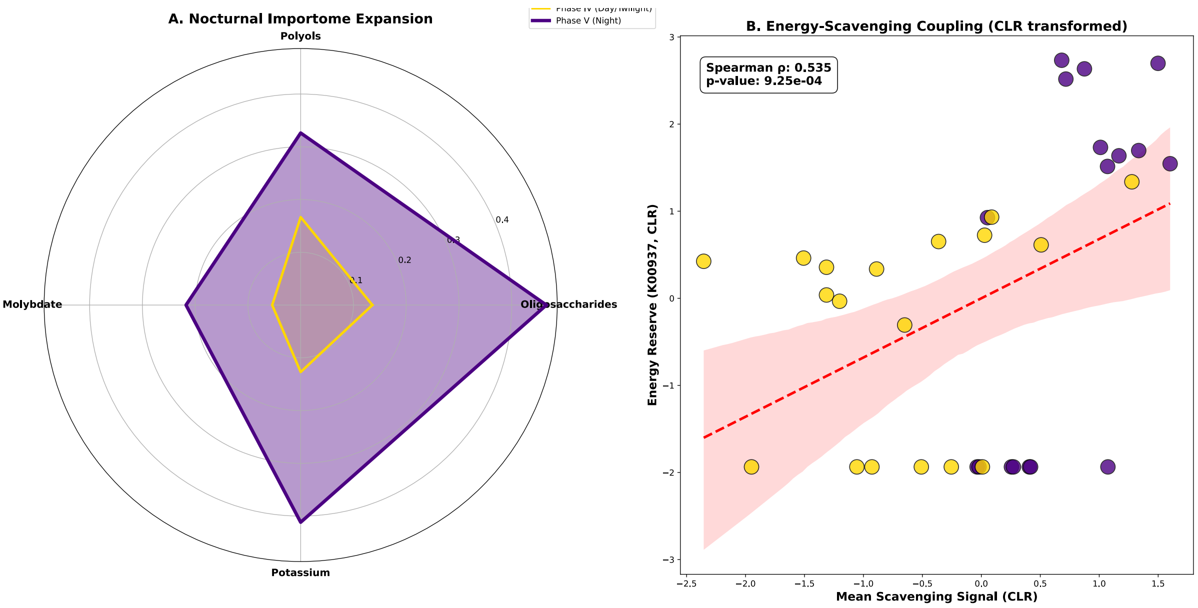


**Figure S8 | Nocturnal importome expansion and energy scavenging coupling.**
**A.** Radar plot illustrating the mean relative transcript abundance of specialized transporters during the day in yellow and during the night in purple. Min-Max normalized values highlight the massive nocturnal induction of carbohydrate, metal, and ion acquisition systems, specifically targeting oligosaccharides (K15770), polyols (K10227), molybdate (K02020), and potassium (K05802). **B** Scatter plot depicting the positive correlation between the integrated scavenging signal, calculated from the mean expression of these key transporters, and cellular energy storage via polyphosphate

kinase (ppk1; K00937). Individual data points numbering 35 represent high-frequency sampling events across the 72-hour time series, color-coded to distinguish daylight samples in yellow from nocturnal samples in purple. Using Centered Log-Ratio transformed transcript counts to eliminate compositional bias, the significant coordination with a Spearman rho of 0.535 and a *p*-value below 0.001 demonstrates a strict metabolic coupling. This relationship confirms that nocturnal nutrient influx directly fuels the replenishment of community energy reserves, effectively bridging the heterotrophic gap before the subsequent dawn activation.

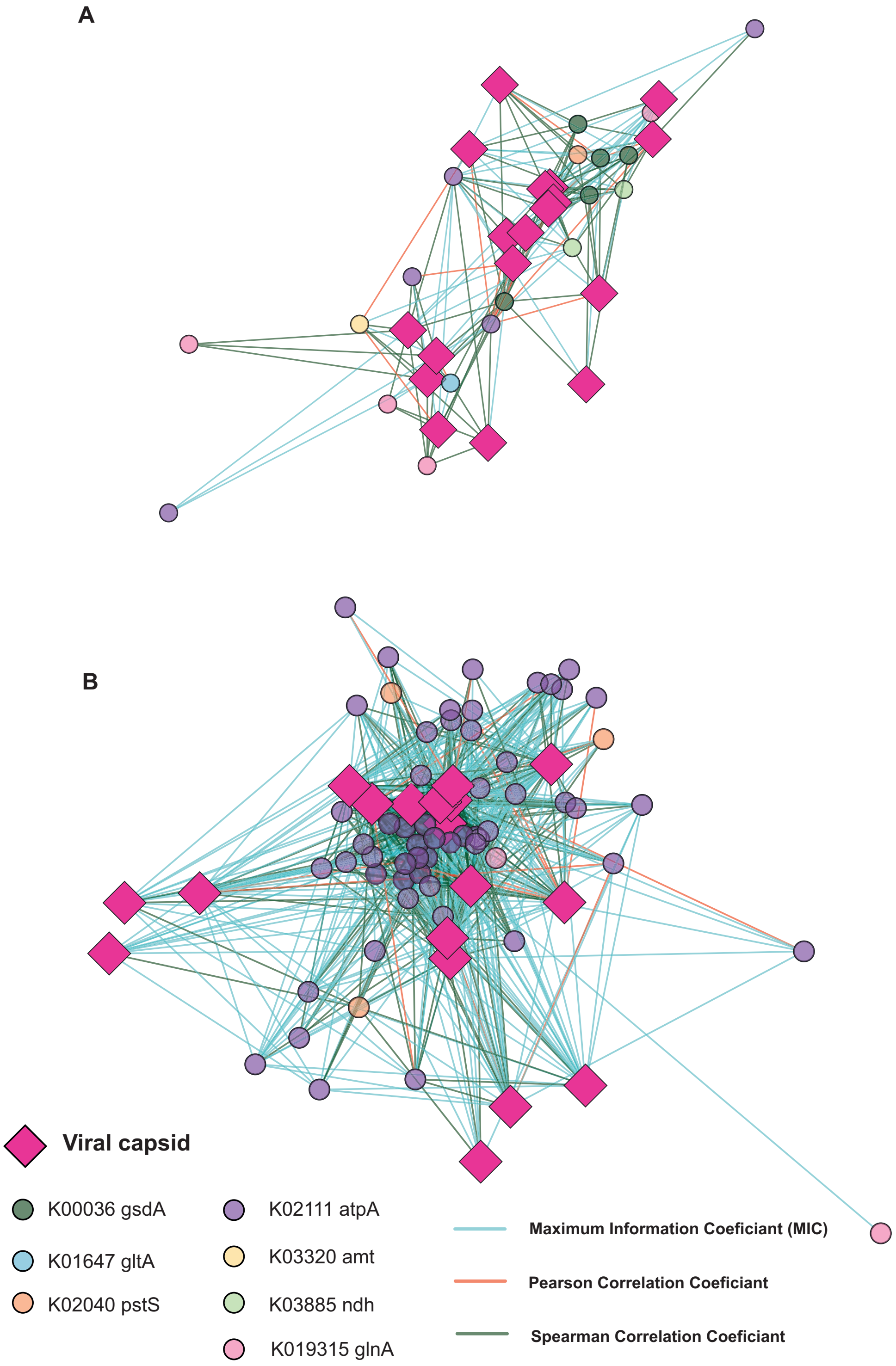

**Figure S9 | Co-expression networks of viral-host metabolic interactions.**

**A.** Positive and **B.** negative correlation networks constructed using a spring embedding layout. Interactions represent significance in at least one of three metrics: Pearson (|*r*| > 0.499), Spearman (|*rho*| > 0.499), or Maximum Information Coefficient (MIC > 0.399). Pink diamonds denote viral capsid proteins; circles indicate host metabolic genes colored by KO function. Edge thickness reflects the maximum correlation coefficient, highlighting the most robust functional dependencies. The network topology illustrates the strategic synchronization of viral assembly with host nitrogen metabolism (**A)** and its antagonism with host energetic maintenance (**B)**.

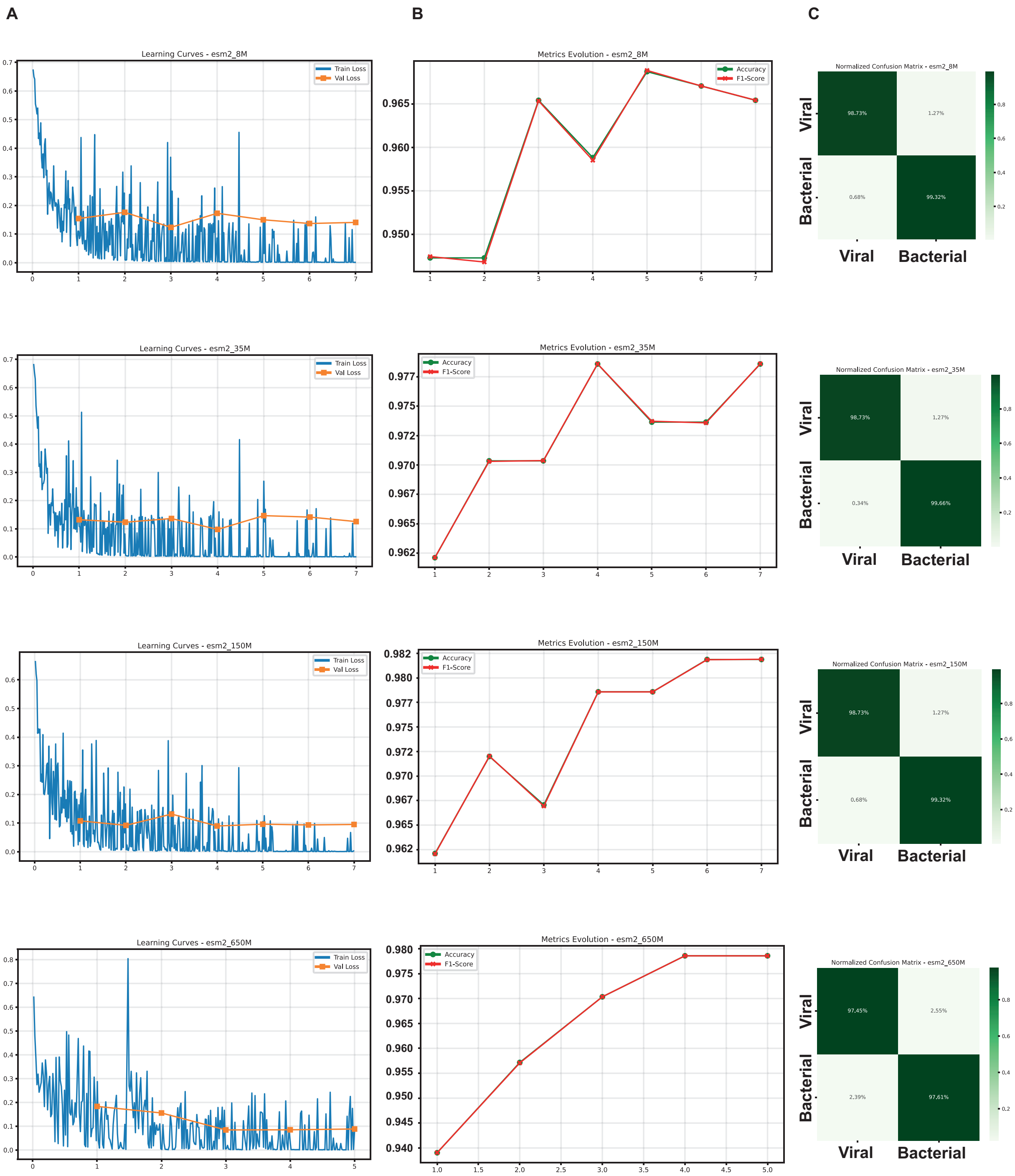


**Figure S10 | Benchmarking and validation of structural classification architectures.**
**A.** Training and validation loss curves across the ESM-2 architecture suite (8 M, 35 M, 150 M, and 650 M parameters). Steady convergence occurs at all scales, with high-parameter models exhibiting

enhanced stability in validation loss. **B.** Longitudinal evolution of classification accuracy and F1-score over training epochs. Performance plateaus are reached earlier in larger architectures, indicating a more efficient latent representation of psbA structural features. **C.** Normalized confusion matrices for the final epoch of each model. Precision remains robust (>97%) across the entire scale range, while high-capacity models (150 M and 650 M) provide the most consistent adjudication between viral and bacterial structural signatures.

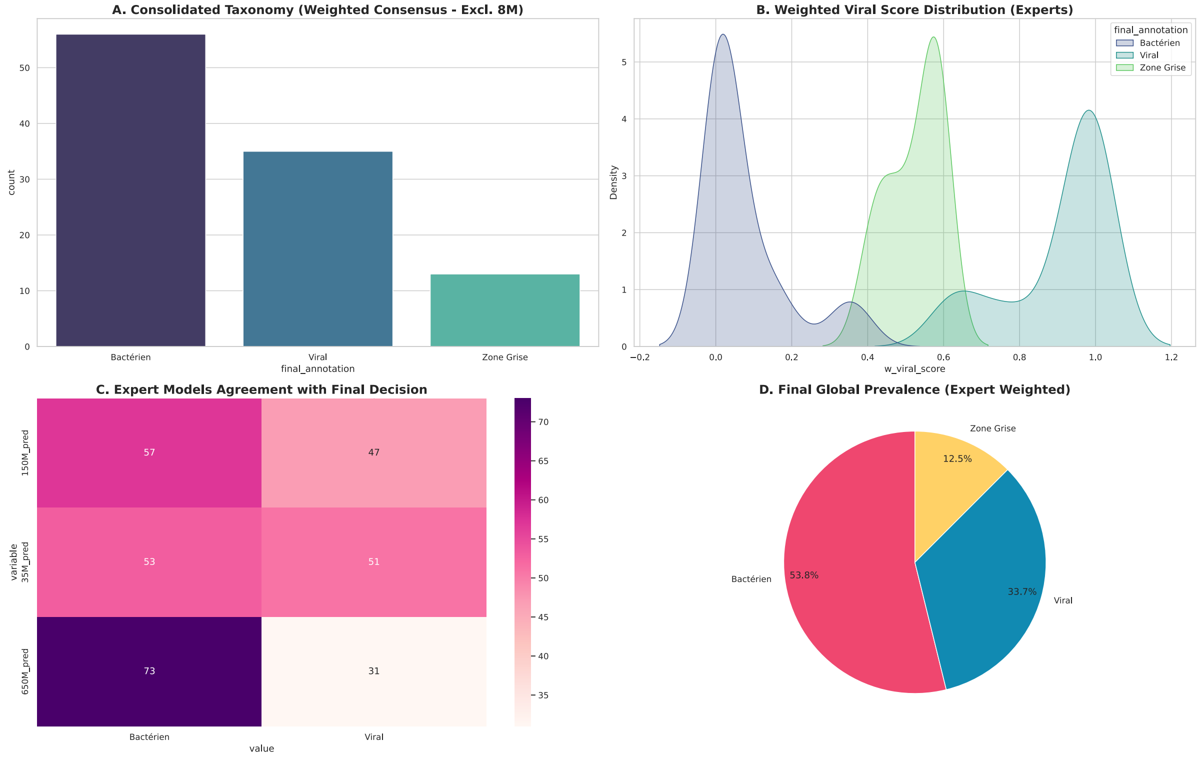


**Figure S11| Structural adjudication of environmental *psbA* transcripts.**
**A.** Consolidated taxonomic distribution of 233 environmental sequences based on a weighted consensus vote across high-capacity ESM-2 models (35M, 150M, and 650M parameters). **B.** Weighted viral score distribution across the expert model ensemble, showing distinct separation between bacterial and viral cohorts with a reduced 'Zone Grise' (12.5%). **C.** Expert models agreement matrix demonstrating the consistency of the 35M, 150M, and 650M architectures in reaching the final weighted consensus annotation. **D.** Final prevalence of confirmed viral *psbA* transcripts (33.7%) compared to bacterial (53.8%) and ambiguous cohorts (12.5%) within the environmental dataset, following weighted multi-model adjudication.

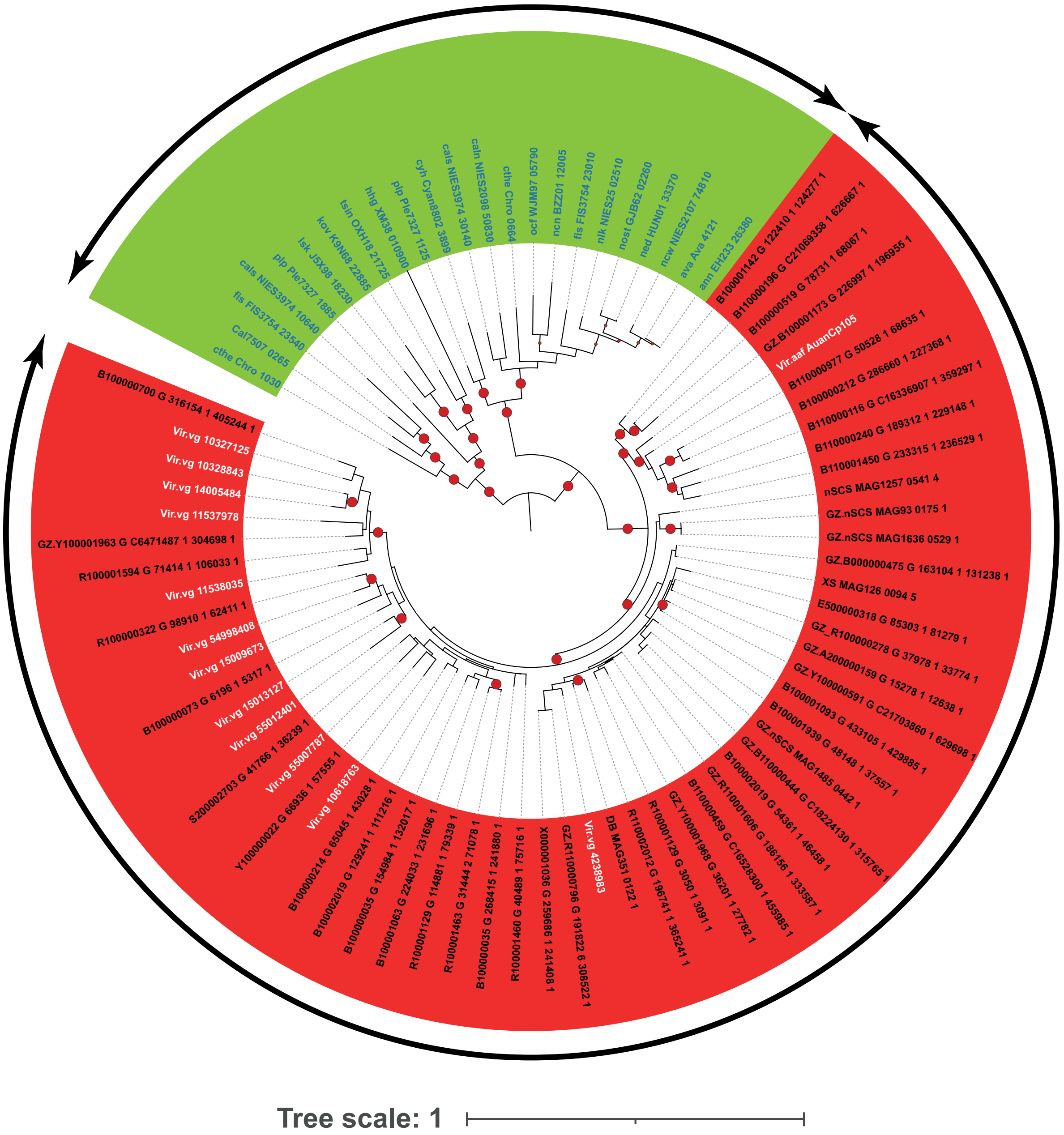


**Figure S12 | Phylogenetic reconstruction of the diel *psbA* pool.**
The maximum likelihood tree was inferred using IQ-TREE with the Q.PFAM+I+R4 substitution model. The tree is rooted using cyanobacterial sequences as the bacterial (Cyano) outgroup (green). Viral query sequences (red) and sequences from the Grey Zone (GZ) are nested within clades of established references (bold), which were sourced from the KEGG database.
